# The capsule of hypervirulent *Klebsiella pneumoniae* maintains neutrophils in neutral activation state

**DOI:** 10.64898/2026.02.24.707717

**Authors:** Karin Santoni, Joshua L.C Wong, Jaie Rattle, Etienne Meunier, Gad Frankel

## Abstract

Hypervirulent *Klebsiella pneumoniae* (hvKp) cause invasive infections despite robust neutrophil recruitment, yet the mechanisms enabling persistence in neutrophil-rich environments remain poorly defined. Here, we show that hvKp do not simply resist neutrophil antimicrobial mechanisms but instead constrain neutrophils into a neutral functional state characterised by limited bactericidal activity. Using genetic dissection of the *rmpADC* locus, pharmacological inhibition of neutrophil effector pathways, and analysis of a diverse panel of clinical isolates, we demonstrate that *rmpADC*-driven capsule properties uncouple neutrophil recognition and activation from bacterial killing. Deletion of *rmpADC* restores phagocytosis, degranulation, and intraphagosomal killing, whereas loss of individual *rmpD* or *rmpC* permits neutrophil activation without bacterial killing. Moreover, hypermucoviscosity alone is sufficient to protect bacteria from neutrophil-mediated killing across multiple genetic backgrounds. Together, these findings identify capsule-driven immune state control as a central mechanism of hvKp neutrophil evasion and reveal distinct thresholds governing neutrophil activation and bactericidal outcome.

## Introduction

*Klebsiella pneumoniae* (Kp) is a Gram-negative pathogen responsible for a wide spectrum of infections, ranging from pneumonia and urinary tract infections to liver abscesses and bacteraemia^1^. Typically, Kp is divided into classical (cKp) and hypervirulent (hvKp) pathotypes^2,3^. cKp are frequently multidrug resistance (MDR) and primarily cause hospital-acquired infections in immunocompromised patients^3–5^. hvKp are distinguished by their ability to cause invasive community-acquired infections in healthy individuals^2–4^. The convergence of cKp and hvKp, leading to MDR hvKp infections in healthy individuals, is an emerging global public health concern^5–8^.

A major virulence factor enhancing pathogenicity in hvKp is the nature of the polysaccharide capsule, the production and physical properties of which are regulated by the *rmpADC* operon, found on either virulence plasmids or the chromosome^9–11^. The *rmp* locus comprises three genes, *rmpA*, *rmpD* and *rmpC*, where RmpA acts a positive regulator of the operon, RmpC drives increased capsule production by upregulating capsule biosynthesis gene expression, and RmpD, which binds the kinase Wzc (a component of the capsule export machinery), induces hypermucoviscosity (HMV)^11–14^. Through these combined effects on capsule, *rmpADC* enhances resistance to complement-mediated killing, antagonises serum bactericidal activity, and impairs phagocytosis, thereby facilitating bacterial persistence, dissemination, and invasive disease in vivo^15–19^. These features are proposed to underpin the ability of hvKp to overcome host immune responses in healthy humans and deletion of any gene in the *rmpADC* operon results in the attenuation in murine pneumonia models^12,14,18^. In the absence of RmpA, D and C cKp lack the typical hvKp capsule features. Nevertheless, RmpD-independent mutations driving HMV in cKp isolates have been identified^20^. In particular, gain-of-function mutations in Wzc that confer constitutive kinase activity, leading to increased capsule polymerisation and chain length, have been identified in clinical cKp isolates^20–22^. These observations indicate that distinct capsule-associated properties have been selected in clinical populations, presumably to benefit the pathogen in shaping host-pathogen interactions.

Neutrophils are the first responders of the innate immune system, rapidly recruited to sites of infection where they coordinate inflammatory signalling and deploy multiple antimicrobial effector functions^23^. In murine models, neutrophil depletion or functional impairment leads to uncontrolled bacterial growth and increased mortality following *Staphylococcus aureus, Acinetobacter baumannii* and *Pseudomonas aeruginosa* infection, underscoring their central role in innate host defence^24–26^. Neutrophils are highly efficient phagocytes, capable of effectively eliminating pathogens^27,28^. Phagocytosis triggers a coordinated antimicrobial program in neutrophils that involves both granule mobilisation and activation of the NADPH oxidase^29^. Upon bacterial engagement and uptake, neutrophil granules containing antimicrobial proteins and proteases, including myeloperoxidase (MPO) and neutrophil elastase (NE), are mobilised and fuse with the forming phagosome, delivering their contents into the intraphagosomal compartment^23,30^. The granule-derived components contribute to bacterial killing through proteolytic degradation of microbial structures and enzymatic activities that impair bacterial viability^23,31,32^. In parallel, phagosomal assembly is accompanied by activation of the NADPH oxidase and production of reactive oxygen species (ROS), which contribute to microbial damage and augment the activity of granule-derived effectors within the phagosome^23,33^. In addition to intracellular deployment, neutrophils can also release granule contents into the extracellular space, particularly under conditions where phagocytosis is limited or incomplete^34^. The extracellular degranulation contributes to antimicrobial defence and inflammatory signalling in the surrounding tissue environment^23,35^. In addition, neutrophils can release extracellular web-like structures, known as neutrophil extracellular traps (NETs), made of decondensed DNA and coated with antimicrobial effectors^36–38^. NETs are released during NETosis, a programmed cell death that depends on ROS production and granule proteins including MPO and NE^39–42^. NETs are particularly effective at immobilising and killing extracellular pathogens^43^. Both MPO and NE play critical roles in host defence against Gramnegative bacteria such as *Pseudomonas aeruginosa* and *E.Coli*, where they contribute to efficient killing^44,45^.

Neutrophil responses are context-dependent, developing along a functional spectrum. They can adopt states ranging from weakly responsive and non-bactericidal to highly inflammatory and microbicidal^46^. Several studies have shown that cKp are efficiently controlled by neutrophils through coordinated antimicrobial mechanisms that include phagocytosis and intracellular killing^44,47^. In contrast, hvKp can persist in vivo alongside neutrophils, suggesting that these strains possess strategies to evade or subvert neutrophil-mediated killing^47–49^. While *rmp*-dependent capsule-mediated resistance to complement and opsonophagocytosis has been extensively described^15,17,50^, how hvKp interact with neutrophils during the earliest stages of infection, and whether they actively modulate neutrophil functional states beyond limiting uptake, remains unexplored.

In this study, we combined genetic dissection of the *rmpADC* locus, pharmacological inhibition of neutrophil effector pathways, and a diverse panel of clinical Kp isolates (spanning multiple capsule and O-antigen backgrounds) to define how hvKp first interacts with neutrophils. We show that hvKp, but not cKp, constrains neutrophils into a neutral functional state characterised by limited effector activation and preserved bacterial survival. We identify the *rmpADC* locus as the central determinant that uncouples Kp neutrophil recognition and activation from bactericidal outcome, revealing distinct thresholds governing neutrophil inflammatory responses and killing. By separating HMV from capsule abundance and extending our findings across multiple genetic backgrounds, this work reframes hvKp immune evasion as a process of immune state control rather than simple resistance to neutrophil phagocytosis and killing.

## Results

### HvKp resist neutrophil-mediated killing

We examined neutrophil responses to hvKp infection using primary bone marrow derived mouse neutrophils, using two hvKp isolates: ICC8001 (our reference strain derived from ATCC43816)^51^ and MRSN16233, a clinical isolate from the MRSN collection^52^. We also included two cKp isolates, MRSN511348 and MRSN740795. All four strains encode K2 capsule and O1αβ,2α LPS O antigen (Table S1). The strains were classified as hvKp if they contained the *rmpADC* operon, displayed HMV, and had a VIR score >1 (indicating the presence of one or more hypervirulence-associated genes), or as cKp if they lacked *rmpADC*, were not HMV, and had VIR scores ≤1^53,54^.

We first quantified bacterial survival following infection of unprimed and LPS-primed neutrophils. Survival was expressed as the percentage of total bacteria recovered in the presence of neutrophils relative to bacteria cultured alone. At multiplicity of infections (MOIs) of 1 and 10, the reference hvKp strain ICC8001 showed 100% survival at 3 and 6 h post-infection (hpi) (Fig. 1A). Similar results were obtained in the presence of either serum or heat-inactivated serum, under LPS-primed and unprimed conditions (Supplementary Figs. S1A, S1B). To determine whether this phenotype extended beyond hvKp ICC8001, we compared survival at 3 hpi in additional hvKp and cKp isolates. While hvKp MRSN16233 exhibited comparable survival to ICC8001, cKp MRSN511348 and MRSN740795 were efficiently killed by neutrophils at both MOIs of 1 and 10 (Fig. 1B). These results indicate that hvKp, but not cKp, resist neutrophil killing.

**Figure 1.**
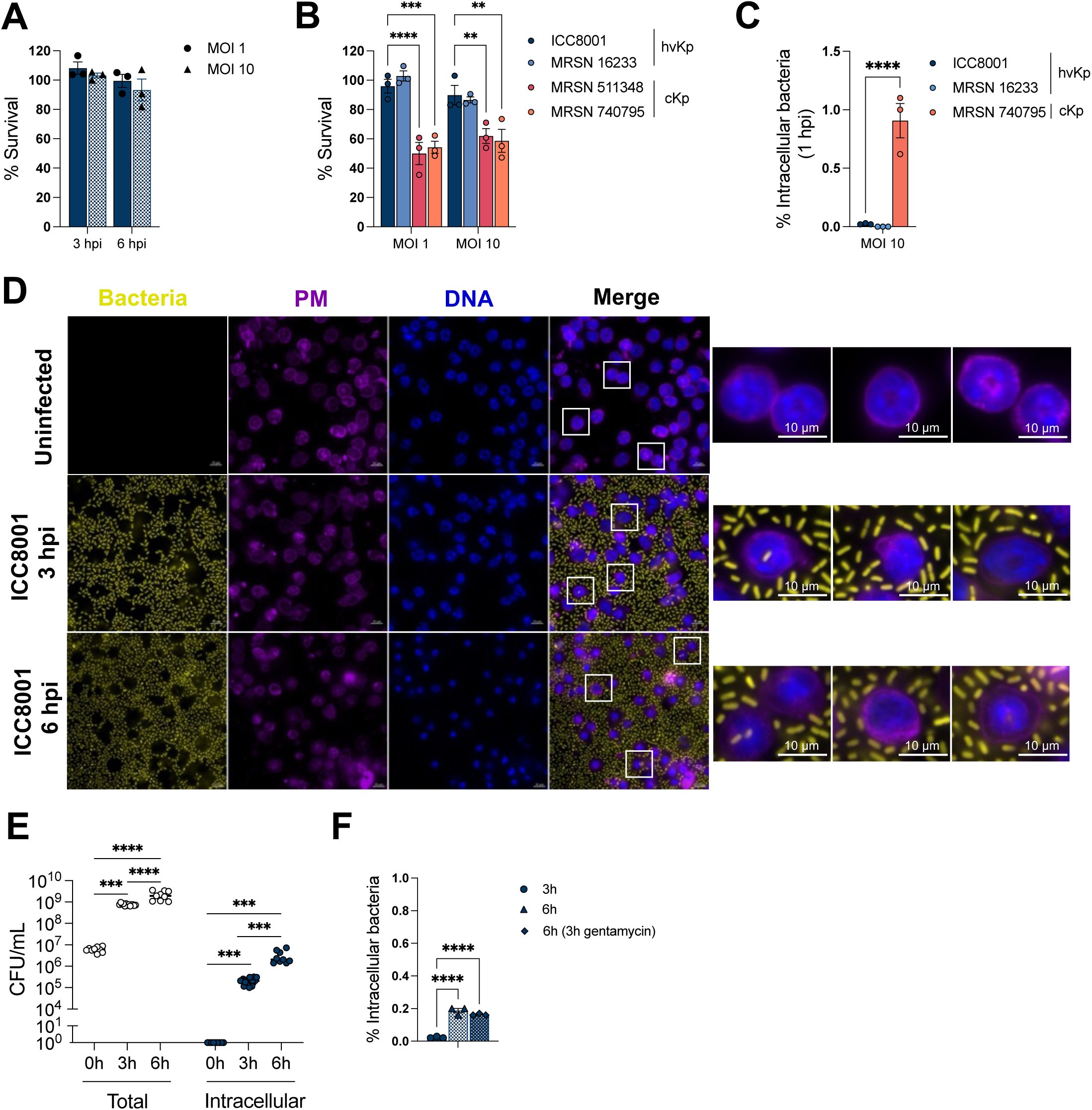
hvKp resist neutrophil-mediated killing. (**A**) Percentage of survival of hvKp ICC8001 in un-primed BMDNs, at 3-6 hpi (MOI 1 and 10). (**B**) Percentage of survival of hvKp and cKp in un-primed BMDNs (MOI 1 and 10), at 3 hpi. (**C**) Percentage of intracellular hvKp and cKp in un-primed BMDNs (MOI 10), at 1 hpi, in the presence of gentamycin. (**D**) hvKp ICC8001 (yellow) localisation in un-primed BMDNs at 3-6 hpi. WGA (purple) and DAPI (blue) were used to stain neutrophils. Scale bars, 10µM. (**E**) Total and intracellular hvKp ICC8001 load in un-primed BMDNs at 0h, 3h and 6 hpi. (**F**) Percentage of intracellular hvKp ICC8001 in un-primed BMDNs (MOI 10), at 3h and 6 hpi, in the presence of gentamycin, added at 3h or 6 hpi. (**A, E**) Significance was determined by ordinary two-way ANOVA, followed by Sidak’s multiple-comparison post-test. (**B**) Significance was determined by ordinary two-way ANOVA compared to ICC8001, followed by Sidak’s multiple-comparison post-test. (**C**) Significance was determined by ordinary one-way ANOVA compared to ICC8001, followed by Dunnett’s multiple-comparison post-test. (**F**) Significance was determined by ordinary one-way ANOVA, followed by Tukey’s multiple-comparison post-test. Values are expressed as mean. **p< 0.01, ***p< 0.001, ****p< 0.0001. Data are representative of at least three independent experiments.

To determine whether hvKp survival reflected evasion of uptake or persistence following phagocytosis, we examined bacterial localisation following infection. As LPS priming and serum did not affect infection outcome, we chose to conduct all the following experiments in serum free conditions. Quantification of intracellular bacteria at 1 hpi using a gentamycin protection assay revealed significantly greater uptake of cKp MRSN740795 compared with hvKp ICC8001 and MRSN16233 (Fig. 1C). The cKp isolate MRSN511348 could not be included in this analysis due to intrinsic resistance to multiple antibiotics, including gentamycin.

As hvKp exhibited minimal uptake at early time points, we next examined later stages of infection to determine whether hvKp is subsequently internalised and, if so, whether it persists intracellularly. Immunofluorescence (IF) microscopy revealed that at 3 and 6 hpi a minority of neutrophils contained intracellular hvKp (Fig. 1D). Enumeration of intracellular CFUs showed that internalised bacteria comprise a small fraction of the total bacterial population, only reaching 0.2% at 6 hpi (Figs. 1E, 1F). To distinguish continuous phagocytosis from primary phagocytosis followed by intracellular replication, we performed gentamicin protection assays. Gentamicin treatment at 3 hpi revealed that intracellular CFU at 6 hpi were comparable to those measured in gentamicin-free conditions, indicating that the intracellular population derives predominantly from early phagocytic events (<3 hpi) followed by intracellular bacterial replication (Fig. 1F).

### HvKp constrains neutrophils into a neutral functional state

We next examined whether the differential survival of hvKp and cKp correlated with differences in neutrophil activation by interrogating major neutrophil effector pathways following infection.

Because NETosis represents a well-described antimicrobial program capable of restricting extracellular pathogens, we first assessed whether Kp triggered NET formation during the early stages of neutrophil interaction. Neither hvKp nor cKp induced neutrophil cell death or NET formation. Assessment of extracellular DNA release using SYTOX^TM^ Green revealed no increase above baseline in any of the Kp strains at early (<6 hpi) or late time points (6-20 hpi), indicating preserved plasma membrane integrity (Supplementary Figs. S1C, 1D). Consistently, IF staining for citrullinated histone H3 (H3cit) and MPO showed no evidence of NET structures at 3 hpi (Fig. 2A). Quantification confirmed the absence of NETosis, with no significant increase in the percentage of H3cit^+^/DAPI^+^ neutrophils or in total H3cit-positive area normalised per 100 neutrophils compared with uninfected controls (Figs. 2B, 2C). These results indicate that, under our infection conditions, NET formation does not contribute to early neutrophil-Kp interactions, consistent with previous reports that NET formation typically requires prolonged stimulation. On this basis and given that hvKp survival diverged from cKp within the first hours of infection, we focused subsequent analyses on effector pathways known to operate rapidly after bacterial engagement, namely ROS production and degranulation.

**Figure 2.**
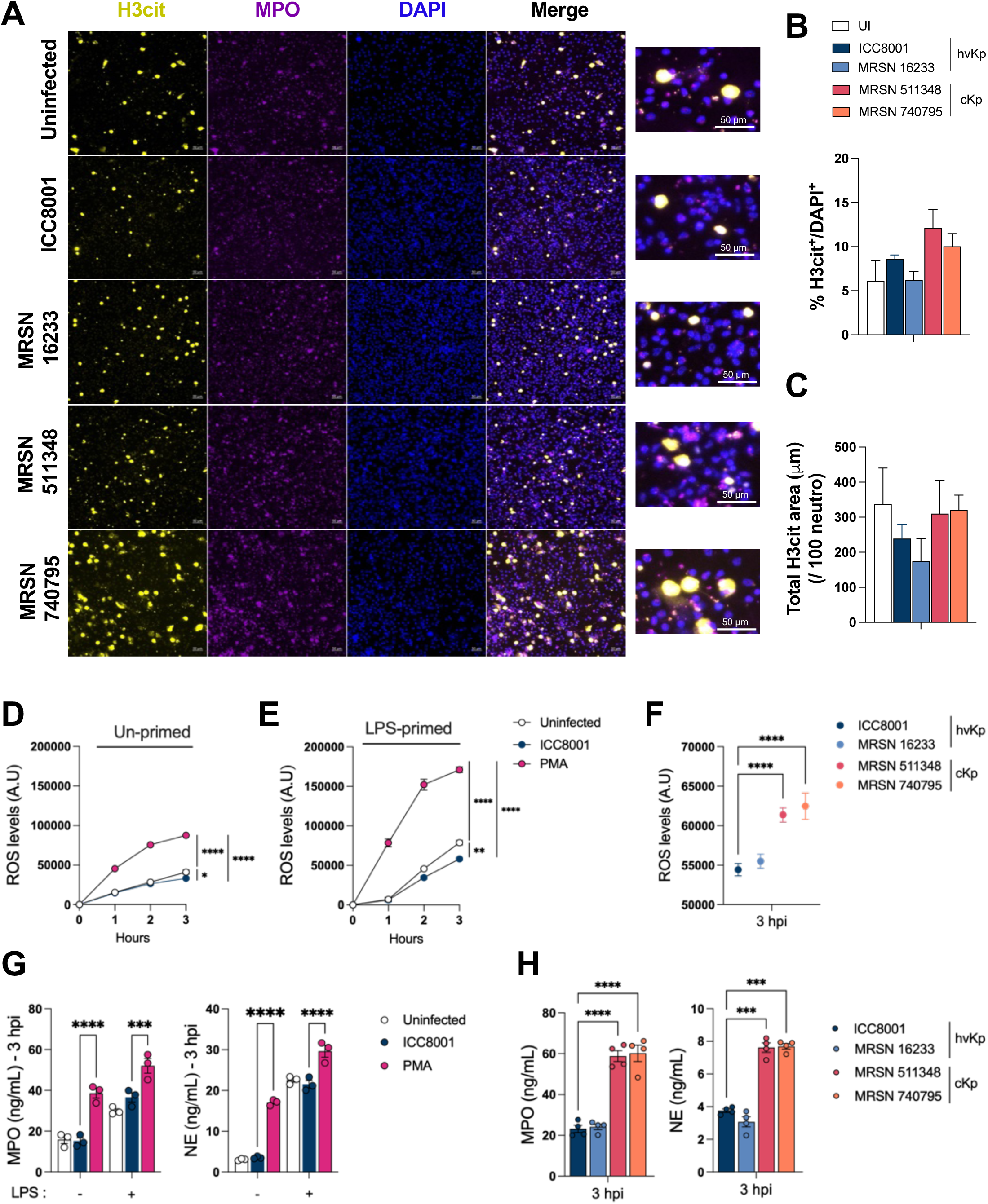
hvKp constrains neutrophils in a neutral functional state. (**A**) Representative immunofluorescence images showing Histone H3 citrullination (H3cit, yellow), MPO (purple) and DAPI (blue) staining of un-primed BMDNs infected with different hvKp and cKp strains for 3h (MOI 10). Scale bars, 50µM. (**B-C**) Quantification of NET formation shown as the percentage of H3cit^+^/DAPI^+^ (**B**) and the total H3cit-positive area normalised per 100 neutrophils (**C**), in uninfected or infected BMDNs with different hvKp and cKp strains for 3h (MOI 10). (**D, E**) ROS levels of BMDNs un-primed (**D**) or LPS-primed (**E**), un-stimulated, stimulated with PMA or infected with hvKp ICC8001 (MOI 10) for 4h. (**F**) ROS levels in un-primed BMDNs, infected with hvKp or cKp for 3 h (MOI 10). (**G**) Levels of NE and MPO in the supernatant of un-primed and LPS-primed BMDNs, un-stimulated, stimulated with PMA or infected with hvKp ICC8001 (MOI 10) for 3 h. (**H**) Levels of NE and MPO in the supernatant of BMDNs, infected for 3 h with different cKp or hvKp (MOI 10). (**D, E**) Significance was determined by ordinary two-way ANOVA, followed by Sidak’s multiple-comparison post-test. (**B-C, F-H**) Significance was determined by ordinary one-way ANOVA compared to ICC8001, followed by Dunnett’s multiple-comparison post-test. Values are expressed as mean. *p< 0.05, **p< 0.01, ****p< 0.0001. Data are representative of at least three independent experiments.

We continuously monitored the production of ROS in neutrophils over 4 h, using the fluorescent probe H₂DCFDA, PMA treatment was used as a positive control. PMA triggered a robust oxidative burst, which was enhanced by LPS priming, validating the assay (Figs. 2D, 2E). In contrast, hvKp did not elicit ROS production above baseline following infection of unprimed and LPS-primed neutrophils (Figs. 2D, 2E). Direct comparison across hvKp and cKp isolates demonstrated that cKp isolates induced significantly increased ROS production compared to hvKp strains at 3 hpi (Fig. 2F).

We next quantified infection-induced neutrophil degranulation. While PMA induced substantial release of MPO and NE, which was enhanced by LPS priming, hvKp ICC8001 triggered minimal degranulation, which was equivalent to the uninfected control neutrophils (Fig. 2G). In contrast, robust MPO and NE release, characteristic of neutrophil responses to bacterial infection *in vitro,* was observed following infection with cKp, compared to hvKp strains (Fig. 2H). As no differences were observed between LPS-primed and unprimed conditions with hvKp, all subsequent experiments were performed using unprimed neutrophils.

Together, these data indicate that neutrophils infected with hvKp enter a neutral, non-inflammatory, state in which early effector responses fail to engage. In contrast, cKp drive an inflammatory neutrophil response that is associated with effective bacterial control.

### Deletion of *rmpADC* restores inflammatory neutrophil responses

A feature that distinguishes hvKp from cKp, is increased production of capsule and HMV, driven by the *rmpADC* operon. However, it is still not known whether the *rmpADC* gene expression is directly implicated in evasion of neutrophil killing. To test this, we compared neutrophil responses to infection with ICC8001 and the isogenic Δ*rmpADC* mutant. The acapsular Δ*wcaJ* mutant was used as a control. Loss of *rmpADC* and *wcaJ* functions were confirmed by absence of HMV and hypercapsulation, as measured by sedimentation assays (OD_600_) and uronic acid quantification (Supplementary Figs. S2A, S2B). In survival assays, the Δ*rmpADC* mutant was efficiently killed by neutrophils at 3 hpi at both MOI of 1 and 10, reaching levels comparable to those observed with Δ*wcaJ* and cKp isolates, whereas wild type ICC8001 was resistant to neutrophil killing (Fig. 3A).

**Figure 3.**
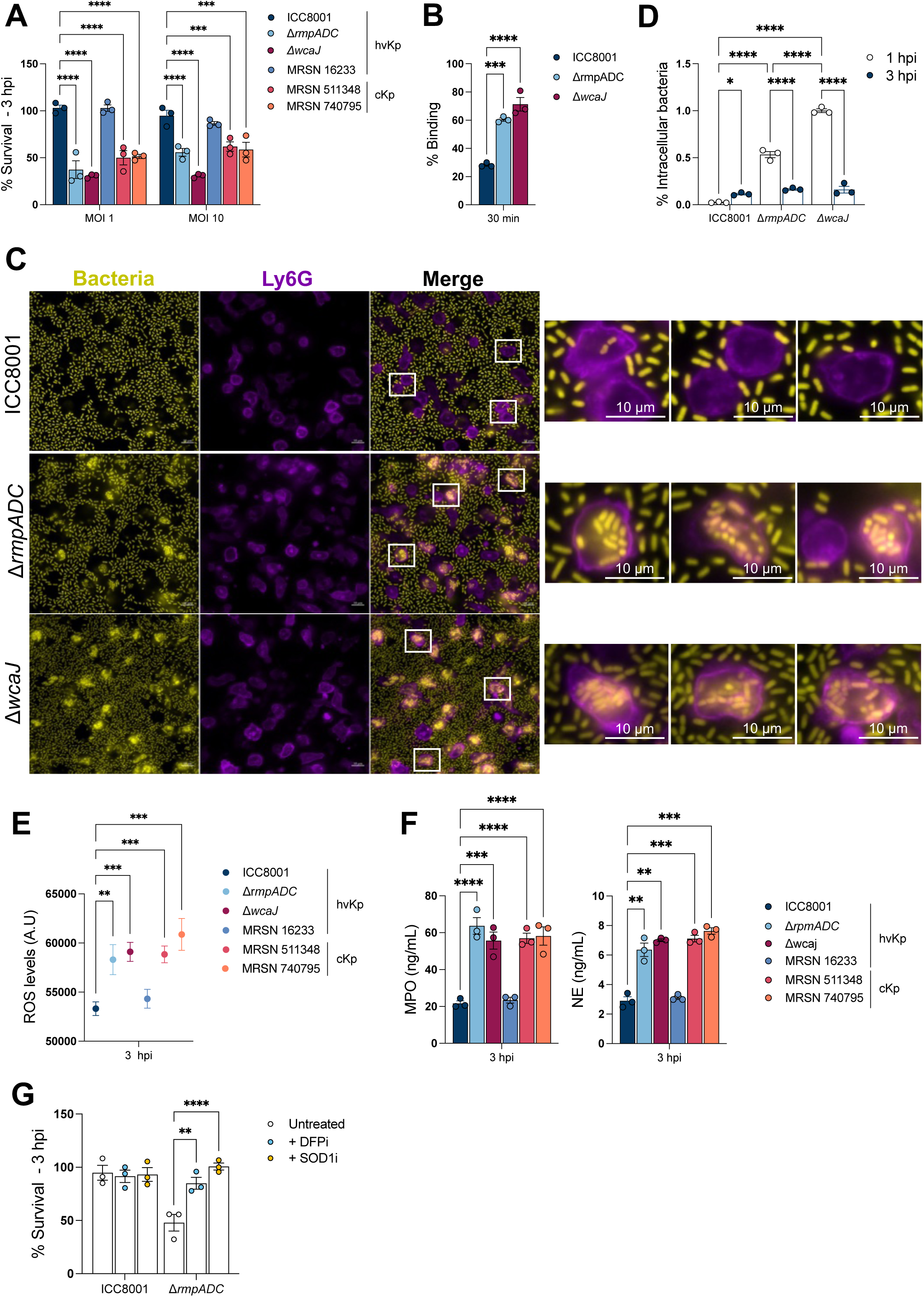
Deletion of *rmpADC* restores inflammatory neutrophil responses in hvKp infection. (**A**) Percentage of survival of different hvKp and cKp in un-primed BMDNs (MOI 1 and 10), at 3 hpi. (**B**) Percentage of binding of hvKp ICC8001, Δ*wcaJ* and Δ*rmpADC* to un-primed BMDNs (MOI 10), at 3 hpi. (**C**) hvKp ICC8001, Δ*wcaJ* and Δ*rmpADC* (yellow) localisation in un-primed BMDNs at 3 hpi. WGA (purple) and DAPI (blue) were used to stain neutrophils. Scale bars, 10µM. (**D**) Percentage of intracellular hvKp ICC8001, Δ*wcaJ* and Δ*rmpADC* in un-primed BMDNs (MOI 10), at 3 hpi. (**E**) ROS levels in un-primed BMDNs, infected with different hvKp or cKp (MOI 10), at 3 hpi. (**F**) Levels of NE and MPO in the supernatant of BMDNs, infected for 3 h with cKp and hvKp (MOI 10). (**G**) Percentage of survival of hvKp ICC8001 and Δ*rmpADC* in un-primed BMDNs, in the presence of DFP and SOD1 (MOI 1), at 3 hpi. (**A**) Significance was determined by ordinary two-way ANOVA compared to ICC8001, followed by Sidak’s multiple-comparison post-test. (**B, E-F**) Significance was determined by ordinary one-way ANOVA compared to ICC8001, followed by Dunnett’s multiple-comparison post-test. (**D, G**) Significance was determined by ordinary two-way ANOVA, followed by Sidak’s multiple-comparison post-test. Values are expressed as mean. **p< 0.01, ***p< 0.001, ****p< 0.0001. Data are representative of at least three independent experiments.

To assess whether the increased susceptibility of the mutants reflected enhanced recognition by neutrophils, we first measured bacterial binding to neutrophils. After 30 min of co-incubation, Δ*rmpADC* mutant associated with neutrophils at significantly higher levels than the parental ICC8001 strain (Fig. 3B), with the acapsular Δ*wcaJ* mutant exhibiting the highest level of association. Microscopy at 3 hpi confirmed that a higher proportion of neutrophils contained intracellular bacteria when infected with Δ*rmpADC* and Δ*wcaJ* compared with ICC8001(Fig. 3C). Quantification of intracellular CFU confirmed enhanced uptake of both mutants at 1 hpi, but intracellular counts declined by 3 hpi to levels comparable to ICC8001 (Fig. 3D), consistent with efficient intracellular killing of the mutant after uptake.

Transmission electron microscopy supported these findings, showing intact hvKp ICC8001 within large intracellular vacuoles at both 3 and 6 hpi, consistent with intracellular bacterial survival (Supplementary Fig. S3A). In contrast, the acapsular Δ*wcaJ* mutant exhibited marked ultrastructural disruption within neutrophils at both 3 and 6 hpi, consistent with intracellular degradation (Supplementary Fig. S3A).

Enhanced recognition of the Δ*rmpADC* mutant was accompanied by restoration of neutrophil effector activation. Deletion of *rmpADC* triggered enhanced ROS production (Fig. 3E) and degranulation of MPO and NE (Fig. 3F), reaching levels comparable to the cKp isolates and Δ*wcaJ* mutant, whereas ICC8001 remained largely non-stimulatory. These results indicate that *rmpADC*-dependent capsule regulation masks hvKp from neutrophil recognition and suppresses effector activation.

To identify the antimicrobial pathways required for killing of the Δ*rmpADC* mutant, we inhibited neutrophil serine proteases with diisopropyl fluorophosphate (DFP), a broadspectrum inhibitor of granule proteases including NE, cathepsin G (CG) and proteinase 3 (PR3), or scavenged ROS using superoxide dismutase 1 (SOD1). Inhibition of either pathway restored Δ*rmpADC* survival to near wild-type levels (Fig. 3G), suggesting that bacterial killing is primarily mediated by protease-dependent killing that requires an intact oxidative burst. This is consistent with previous observations that intraphagosomal protease activity depends on NADPH-oxidase-derived ROS, required for optimal protease activation within phagosomes^32^. Thus, inhibition of ROS generation production is likely to indirectly impair protease-mediated killing. Neither inhibitor impacted on ICC8001 survival.

Together, these data indicate that expression of *rmpADC* shields hvKp from neutrophil recognition and maintains neutrophils in a neutral, non-inflammatory state. The *rmpADC* mutant and cKp strains induce inflammatory effector activation, similarly to the acapsular mutant, and susceptibility to intraphagosomal killing is driven primarily by granule-derived serine protease.

### Individual *rmpADC* components differentially regulate neutrophil activation and killing

To determine the contribution of individual genes within the *rmpADC* operon, we compared the parental hvKp ICC8001 with isogenic mutants lacking *rmpA*, *rmpD* or *rmpC* individually, alongside the Δ*rmpADC* deletion mutant. As these genes lie in an operon, we introduced premature stop codons (*rmpA* STOP*, *rmpD* STOP* and *rmpC* STOP*) into each gene to preclude polar effects induced by large open read frame deletions. Functional validation of the single *rmp STOP* mutants confirmed phenotypes consistent with their known roles in capsule regulation, with *rmpA*STOP* phenocopying deletion of the full operon, *rmpD*STOP* abolishing HMV and *rmpC*STOP* selectively reducing capsule abundance (Supplementary Figs. S2C, S2D). Survival assays revealed two distinct phenotypes. Deletion of *rmpA*, as observed with the Δ*rmpADC* deletion (Fig. 3A), resulted in efficient neutrophil-mediated killing (Fig. 4A). This finding is in keeping with the known regulatory effect of RmpA as the primary positive autoregulator of the operon. In contrast, loss of *rmpD* or *rmpC* independently did not trigger neutrophil killing with either of the single mutants maintaining high survival indistinguishable from ICC8001 (Fig. 4A). These data indicate that either HMV (RmpD) or increased capsulation (RmpC) are sufficient, independently, to resist killing by neutrophils.

**Figure 4.**
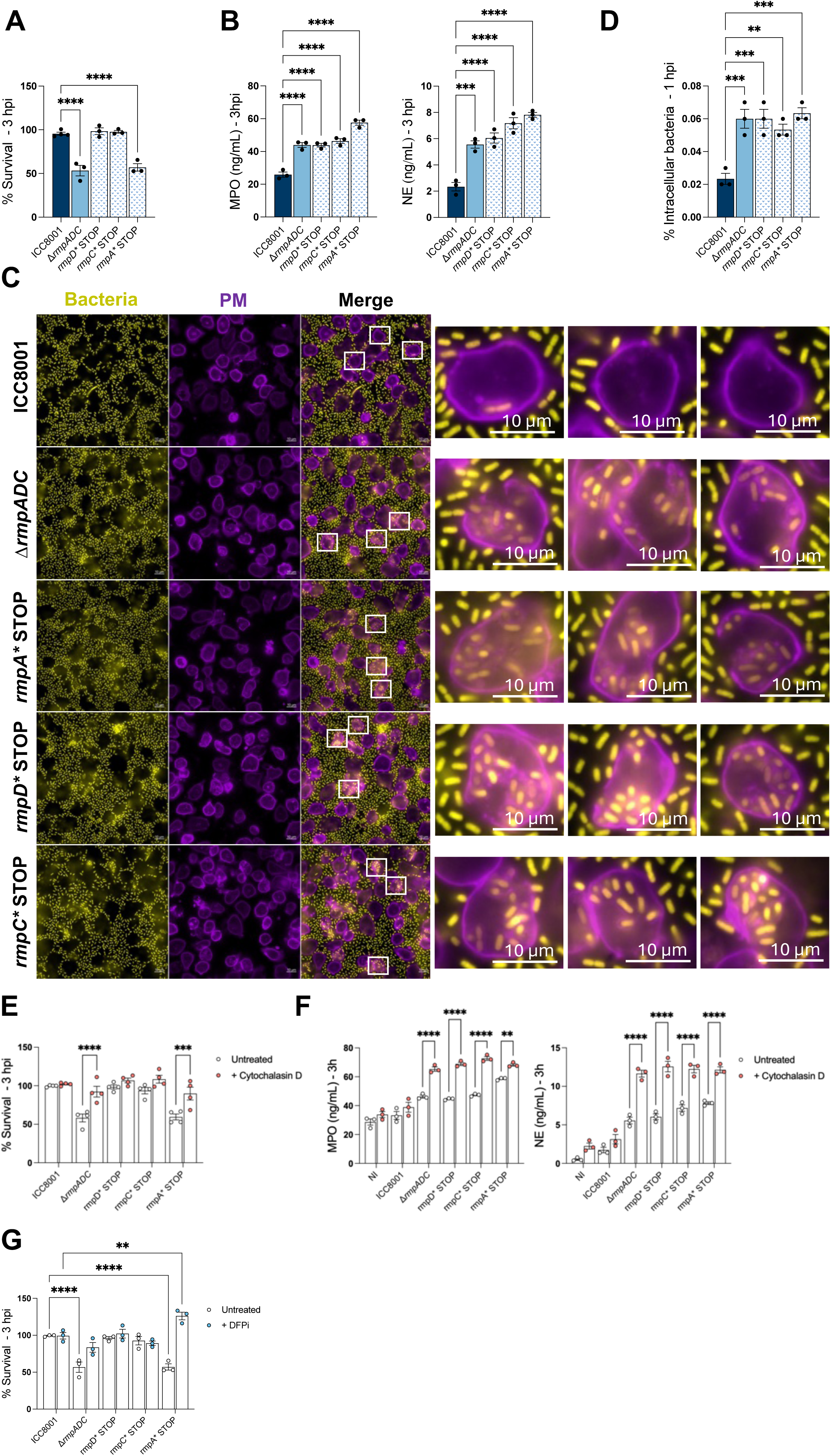
Individual *rmpADC* genes differentially regulate neutrophil activation and killing. (**A**) Percentage of survival of hvKp ICC8001 and isogenic strains in unprimed BMDNs, (MOI 1), at 3 hpi. (**B**) Levels of NE and MPO in the supernatant of unprimed BMDNs, infected for 3 h with hvKp ICC8001 and isogenic strains (MOI 10). (**C**) hvKp ICC8001 and isogenic strains (yellow) localisation in un-primed BMDNs at 3 hpi. WGA (purple) and DAPI (blue) were used to stain neutrophils. Scale bars, 10µM. (**D**) Percentage of intracellular hvKp ICC8001 and isogenic strains in un-primed BMDNs (MOI 10), at 1 hpi. (**E**) Percentage of survival of hvKp ICC8001 and isogenic strains in un-primed BMDNs, in the presence of Cytochalasin D (MOI 1), at 3 hpi. (**F**) Levels of NE and MPO in the supernatant of un-primed BMDNs, infected for 3 h with hvKp ICC8001 and isogenic strains (MOI 10), in the presence of Cytochalasin D. (**G**) Percentage of survival of hvKp ICC8001 and isogenic strains in un-primed BMDNs, in the presence of DFP (MOI 1), at 3 hpi. (**A-B, D**) Significance was determined by ordinary one-way ANOVA compared to ICC8001, followed by Dunnett’s multiple-comparison post-test. (**E-G**) Significance was determined by ordinary two-way ANOVA, followed by Sidak’s multiple-comparison post-test. Values are expressed as mean. **p< 0.01, ***p< 0.001, ****p< 0.0001. Data are representative of at least three independent experiments.

We next examined if survival of the single mutants was associated with suppression of neutrophil effector responses. Despite their resistance to killing, *rmpD*STOP* and *rmpC*STOP*, as well as *rmpA*STOP*, induced robust degranulation, with MPO and NE release significantly increased relative to ICC8001 (Fig. 4B). Consistent with neutrophil activation, microscopy showed increased internalisation compared to ICC8001 (Fig. 4C). Intracellular CFU quantification confirmed significant enhanced mutant uptake early after infection (1 hpi) (Fig. 4D). These results suggest that loss of either HMV (RmpD) or increased capsulation (RmpC) is sufficient to restore neutrophil recognition, phagocytosis, and degranulation. However, despite appropriate neutrophil effector functions, HMV or increased capsulation is independently sufficient to protect Kp from killing by neutrophils.

Because neutrophil activation was restored in the *rmpD*STOP* and *rmpC*STOP* mutants without resulting in bacterial killing, we next asked whether the spatial deployment of antimicrobial effectors, rather than their activation per se, determines bactericidal outcome. Specifically, we tested whether effective killing requires delivery of granule contents into the phagosome rather than their release extracellularly. To address this, we inhibited actin-dependent phagocytosis using cytochalasin D, an approach that allows discrimination between killing mechanisms dependent on intraphagosomal delivery of granule contents and those mediated extracellularly. Blocking phagocytosis selectively increased survival of the Δ*rmpADC* and *rmpA*\*STOP mutants (Fig. 4E), demonstrating that killing of these strains requires internalisation and subsequent intraphagosomal deployment of antimicrobial effectors. In contrast, cytochalasin D had no effect on survival of the *rmpD*\*STOP and *rmpC*\*STOP single mutants (Fig. 4E), consistent with their resistance to killing even when phagocytosed.

Analysis of degranulation under these conditions revealed that inhibition of phagocytosis did not impair granule mobilisation but altered its spatial deployment. Specifically, cytochalasin D treatment resulted in increased extracellular release of MPO and NE during infection with all *rmp* mutants, but not with ICC8001 (Fig. 4F), indicating that this effect is restricted to strains that are recognised and activate neutrophils. Together, these data demonstrate that phagocytosis determines whether neutrophil granule contents are delivered into phagosomes or released extracellularly.

Having established that effective killing requires intraphagosomal delivery of granule contents, we next asked whether the differential survival of *rmp* locus mutants reflected altered sensitivity to intraphagosomal granule proteases. To test this, neutrophils were treated with the serine protease inhibitor DFP before infection with the isogenic mutant strains. Protease inhibition selectively increased survival of Δ*rmpADC* and *rmpA*STOP* mutants but had no effect on *rmpD*STOP* and *rmpC*STOP* mutants (Fig. 4G), despite their ability to induce phagocytosis and degranulation.

Together, these findings demonstrate that neutrophil-mediated killing of hvKp requires both delivery of granule contents into the phagosome and loss of *rmpD*- and *rmpC*-dependent protection. While partial disruption of the rmp system restores neutrophil activation and uptake, expression of either RmpD or RmpC is sufficient to protect bacteria from protease-dependent killing within the phagosome.

### Capsule-driven HMV promotes a neutral state across hvKp lineages

To determine whether the *rmpADC*-dependent neutrophil phenotype observed in ICC8001 is conserved across the Kp species complex, we analysed a panel of clinical isolates spanning diverse K and O antigen backgrounds. This panel included hvKp strains carrying *rmpADC* as well as cKp strains lacking *rmpADC*. Across this panel, bacterial survival following neutrophil co-incubation closely correlated with the presence of *rmpADC*. Strains carrying *rmpADC* were consistently resistant to neutrophil-mediated killing, whereas *rmpADC*-negative isolates were efficiently killed (Fig. 5A). Degranulation across these isolates followed the expected inverse pattern, with *rmpADC*-positive strains eliciting lower MPO and NE release compared to *rmpADC*-negative strains (Fig. 5B), consistent with the neutrophil “neutral state” associated with hvKp infection. Importantly, this expands our findings beyond K2 capsule as K3, K10, K19, K20, K51 and K64 were included in this analysis, demonstrating that *rmpADC*-associated neutrophil phenotype is conserved across diverse Kp capsule types.

**Figure 5.**
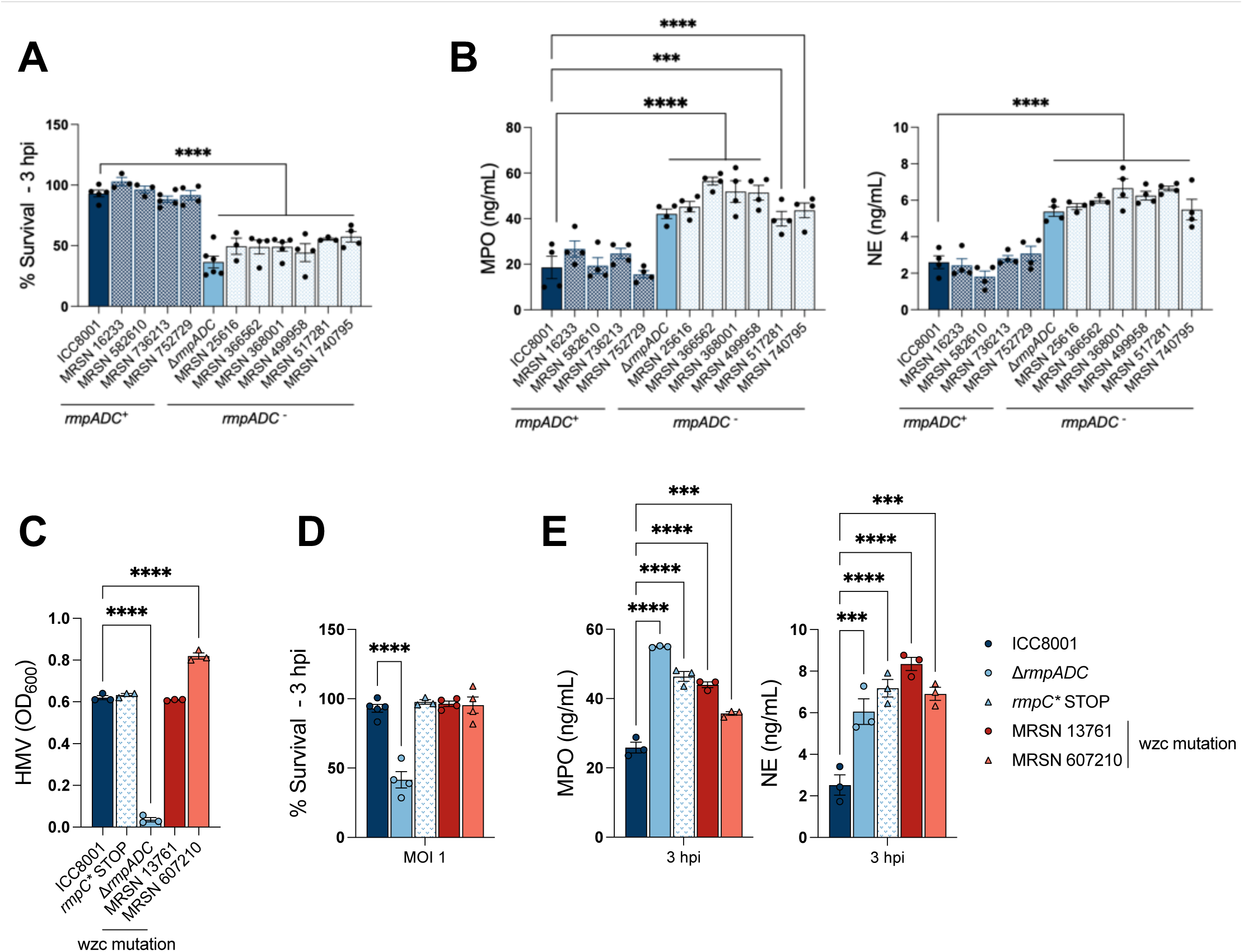
Capsule-driven hypermucoviscosity promotes a neutral sate across hvKp lineages. (**A**) Percentage of survival of hvKp and cKp in un-primed BMDNs (MOI 1), at 3 hpi. (**B**) Levels of NE and MPO in the supernatant of un-primed BMDNs, with hvKp and cKp (MOI 10), at 3 hpi. (**C**) HMV of hvKp ICC8001, isogenic strains and cKp carrying the *wzc* mutation, expressed as OD_600_ of the culture supernatant. (**D**) Percentage of survival of hvKp ICC8001, isogenic strains and cKp carrying the *wzc* mutation, in un-primed BMDNs (MOI 1), at 3 hpi. (**E**) Levels of NE and MPO in the supernatant of un-primed BMDNs, infected for 3 h with hvKp ICC8001, isogenic strains and different cKp carrying the *wzc* mutation (MOI 10). (**A-E**) Significance was determined by ordinary one-way ANOVA compared to ICC8001, followed by Dunnett’s multiple-comparison post-test. Values are expressed as mean. **p< 0.01, ***p< 0.001, ****p< 0.0001. Data are representative of at least three independent experiments.

We then identified cKp isolates (MRSN13761, MRSN607210) harbouring mutations in *wzc*, which confer an RmpD-independent HMV phenotype (Fig. 5C), allowing us to test whether this physical capsule feature alone is sufficient recapitulate hvKp neutrophil response in *rmpADC-*negative cKp isolates. These strains, harbouring naturally evolved *wzc* mutations, were found to be resistant to neutrophil killing (Fig. 5D) with no observable difference between survival of these strains and hvKp ICC8001. However, unlike *rmpADC*-positive hvKp, these HMV *wzc* mutants still triggered robust degranulation, with elevated MPO and NE release (Fig. 5E). This phenotype closely mirrors that observed with the ICC8001 *rmpD*STOP* and *rmpC*STOP* single mutants, which also induced strong neutrophil degranulation while resisting neutrophil killing. Together, these findings indicate that HMV alone allows bacteria to persist despite neutrophil recognition and degranulation, maintaining neutrophils in a functionally inflammatory but non-bactericidal state rather than driving an effective inflammatory response.

## Discussion

HvKp causes invasive infections despite encountering functional first responder neutrophils. While capsule-associated immune evasion has long been recognised as a hallmark of hvKp, with prior work largely focused on capsule-mediated resistance to opsonophagocytosis or serum killing^15,17,50^, how the capsule shapes neutrophil functional states and evades neutrophil-mediated clearance has remained incompletely understood. Here, we demonstrate that hvKp do not merely resist neutrophil antimicrobial mechanisms, but instead actively constrains neutrophils in a neutral functional state characterised by limited effector activation, uptake, and bacterial killing. Using isogenic mutants, pharmacological inhibition, and a diverse panel of clinical isolates, we identify *rmpADC*-driven hypermucoviscosity as a central determinant that uncouples neutrophil recognition and activation from bactericidal outcome.

Our initial experiments establish a clear functional dichotomy between hvKp and cKp. While cKp rapidly triggered ROS production, degranulation, and bacterial killing, hvKp persisted despite prolonged neutrophil contact. Importantly, this persistence was not explained by delayed kinetics alone: hvKp failed to induce early ROS or granule release, and only a small fraction of bacteria were internalised, even at later time points. These findings define a baseline neutral neutrophil state during hvKp infection, in which antimicrobial programs remain largely disengaged.

Genetic dissection of hvKp revealed that this neutral state is actively enforced by the *rmpADC* operon. Deletion of *rmpADC* restored neutrophil binding, uptake, effector activation, and killing, phenocopying complete capsule loss and demonstrating that *rmpADC* is necessary and sufficient to mask hvKp from neutrophil engagement. Mechanistic analysis revealed that killing of the Δ*rmpADC* mutant required intact intraphagosomal antimicrobial activity, as inhibition of either serine proteases or ROS was sufficient to abrogate killing. The comparable effects of DFP and SOD1 suggest that protease-dependent killing is central in this context, with oxidative pathways likely supporting optimal protease function rather than acting as the dominant bactericidal mechanism. In contrast, killing of acapsular bacteria was selectively ROS-dependent, highlighting distinct effector requirements depending on capsule architecture.

A key conceptual insight emerged from analysis of individual *rmpADC* components. While deletion of *rmpA* restored killing, loss of *rmpD* or *rmpC* alone uncoupled neutrophil activation from killing: these mutants were readily recognised, phagocytosed, and induced robust degranulation, yet remained resistant to killing. Cytochalasin D and protease inhibition experiments demonstrated that phagocytosis and granule delivery occurred but were insufficient to drive bactericidal activity when either *rmpD* or *rmpC* was expressed. These findings reveal distinct thresholds governing neutrophil responses: recognition and activation versus effective killing, and show that hvKp prevent neutrophils from transitioning from a neutral state into a productive inflammatory state.

Importantly, this framework extends beyond a single strain or lineage. Analysis of a diverse panel of clinical isolates demonstrated that resistance to neutrophil killing is strongly linked with hypermucoviscosity rather than capsule serotype or genetic background. cKp strains rendered hypermucoviscous through *wzc* mutation were highly resistant to killing despite inducing strong degranulation, closely mirroring the phenotype of *rmpD*- and *rmpC*-deficient ICC8001. These findings demonstrate that hypermucoviscosity alone is sufficient to protect bacteria downstream of neutrophil recognition, even in the absence of *rmpADC*-driven suppression of activation. This observation provides critical insight into how capsule architecture and physical properties can modulate immune outcomes independently of canonical inflammatory signalling.

Our findings also argue for caution in defining hypervirulence solely based on *rmpADC* expression. cKp isolates rendered hypermucoviscous through *wzc* mutation displayed hvKp-like resistance to neutrophil killing despite lacking *rmpADC* and other canonical hypervirulence markers. These data indicate that functional immune evasion phenotypes, rather than genetic markers alone, may better predict neutrophil susceptibility and pathogenic potential. In this context, hypermucoviscosity emerges as a property capable of conferring hvKp-like behaviour even in a classical genetic background, underscoring the need to consider capsule architecture and physical properties when assessing virulence.

Collectively, our results redefine hvKp–neutrophil interactions as an active process of immune state control rather than passive resistance. HvKp enforce a neutral neutrophil state that can tolerate recognition and even partial activation without progressing to bactericidal inflammation. This strategy likely provides a selective advantage during early infection, allowing hvKp to persist in neutrophil-rich environments. More broadly, these findings suggest that targeting bacterial determinants of intraphagosomal resistance, rather than amplifying neutrophil activation, may represent a more effective strategy to restore neutrophil-mediated clearance of hvKp.

## Acknowledgments

We would like to thank the METi imaging facility (CBI, Toulouse), member of the national infrastructure France-BioImaging supported by French National Research Agency (ANR-10-INBS-04) for the Transmission Electron Microscopy images. This project is supported by a Wellcome Trust Investigator Award grant 224282/Z/21/Z.

## Author contributions

K.S performed all experiments and generated the single STOP codon mutants *rmpA*- D*-C* STOP*. J.R generated the Δ*rmpADC* mutant. W.W.L generated the Δ*wcaJ* mutant. E.M provided advice and contributed to TEM experiments. K.S and G.F wrote the manuscript. J.W. provided advice on the manuscript. G.F directed the study.

## Materials and Methods

### Generation of hvKp mutants

All genomic mutations were made in ICC8001, a rifampicin-resistant derivative of *K. pneumoniae* ATCC43816 using a two-step recombination methodology, as described^55^. Strains used are listed in supplementary table 1.

### Bacterial Cultures

*K. pneumoniae* (Kp) strains were grown overnight in LB medium, containing 50µg/mL of streptomycin, at 37°C with constant agitation (200 rpm), and sub-cultured the next day by diluting overnight culture 1:100 at 37°C, with constant agitation (200 rpm), until reaching an optical density (OD) of 1. All *K. pneumoniae* strains used in the study are listed in supplementary table 1.

### Isolation of primary murine neutrophils

Murine bone marrow cells were isolated from tibias and femurs of C57BL/6 mice. Neutrophils were purified by positive selection using Anti-Ly-6G MicroBead Kit (Miltenyi Biotech) according to manufacturer’s instructions. This process routinely yielded >95% of neutrophil population as assessed by flow cytometry of Ly6G^+^/CD11b^+^ cells.

### Cell plating and infection of neutrophils

Following isolation, neutrophils were centrifugated for 10 min at 300xg and pellet was resuspended in serum free Opti-MEM medium (Gibco). Absolute cell number was determined with automated countess 3 cell counter (Invitrogen) with trypan blue cell death exclusion method (typically living cells represent > 80% of cell solution) and cell density was adjusted at 10^6^ / mL by adding Opti-MEM culture medium. Neutrophils were then plated in either 96 well plates, 24 well plates or 6 well plates with 50 µL (5.10^4^ cells), 500 µL (5.10^5^ cells) or 2 mL (2.10^6^ cells) respectively. When indicated, cells were primed with LPS (200 ng/ml) for 2 hours. Neutrophils were infected with various bacterial strains and multiplicity of infections (MOI) as indicated in figure legends, centrifuged for 5 min at 700xg and incubated at 37°C in a 5% carbon dioxide (CO_2_) incubator the indicated time.

### ELISA(s)

Mouse NE (R&D, DY4517) and MPO (R&D, DY3667) levels were quantified according to the manufacturer’s instructions.

### Quantification of ROS levels

Cells were plated at density of 5.10^4^ cells per well in Black/Clear 96-well Plates (Nunc) in Opti-MEM culture medium supplemented with the cell-permeant 2’,7’-dichlorodihydrofluorescein diacetate (H2DCFDA) (10µM) (ThermoFisher, D399) and infected/treated as mentioned in figure legends. Fluorescence was measured in realtime using FLUOstar Omega plate reader (BMG Labtech) at 37°C with 5% CO_2_.

### Quantification of CFU loads / survival

Neutrophils were plated at a density of 5.10^5^ cells per well in a 24-well plate in Opti-MEM culture medium and infected with different Kp strains at an MOI of 1 or 10 for 3 to 6h. At the end of the infection, total bacterial load was determined by lysing cells with 0.1% Triton X-100 for 10 min and subjected to 10-fold serial-dilution before plating on agar plates to enumerate colony-forming units (CFU) and incubated overnight at 37°C. Bacterial survival was expressed as the ratio of total CFU recovered from neutrophil-infected wells relative to CFU recovered from bacteria cultured in parallel in the absence of neutrophils for the same duration (3-6 hpi), thereby normalizing for strainspecific growth.

Phagocytosis assay was performed by adding gentamycin (100µg/mL) for 30min, at either 30min or 2h post-infection to eliminate extracellular bacteria, washing 3 times the cells before adding fresh Opti-MEM medium with 0.1% Triton X-100. Intracellular CFU counts was determined by performing a 10-fold serial-dilution before plating on agar plates and incubated overnight at 37°C. Intracellular bacterial counts were expressed relative either to CFU recovered from bacteria cultured alone for the same duration (3-6 hpi), or for early uptake measurements (1 hpi) relative to the initial inoculum (T_0_), allowing discrimination between bacterial uptake and subsequent intracellular killing.

### Immunofluorescence

Neutrophils were plated at a density of 5.10^5^ cells on glass bottom Ibidi chambered coverslips in Opti-MEM culture medium and infected at an MOI of 10 for 3 to 6h. At the end of the infection, cell supernatant was removed, and cells were fixed with a 4% PFA solution for 10 min at 37°C. When specified, plasma membrane was stained with Wheat Germ Agglutinin-Alexa Fluor 647 (WGA) (ThermoFisher, W32466) at 1:250 dilution in HBSS and incubated for 10 min in the dark. Permeabilization was performed by incubating cells for 10 min in PBS containing 0.1% Triton X-100. To block unspecific binding of antibodies, cells were incubated in PBS-T (PBS + 0.1% Tween 20), containing 2% BSA, for 30 min. Cells were incubated with primary antibodies (1:100) anti-MPO (abcam, ab300650), anti-H3cit (abcam, ab5103), overnight at 4°C in PBS-T + 2% BSA solution and detected with the appropriate secondary antibodies (1:1000) Alexa Fluor 555-conjugated donkey anti-rat (ThermoFisher, A21434) or AlexFluor 488-conjugated goat anti-rabbit (ThermoFisher, A21206), for 1 hour at room temperature. DNA was counterstained with DAPI (1:1000). Stained cells were imaged using Zeiss AxioVision Z1 microscope and analysed with Zen 2.3 Blue Version (Carl Zeiss MicroImaging GmbH, Germany).

### Quantification of NETs

NET formation was quantified using two complementary metrics: (i) the percentage of H3cit-postitive cells and (ii) total H3cit-positive area normalised to cell number.

i. Neutrophils were identified by DAPI-positive nuclei. H3cit positivity was determined by applying a consistent intensity threshold to the H3cit channel across all conditions within a given experiment. The proportion of H3cit^+^/DAPI^+^ neutrophils was calculated per field and averaged per condition.
ii. The H3cit-positive area was segmented using a fixed threshold and measured as total area (μm^2^) per field. Values were normalised to the number of DAPI-positive neutrophils in the same field and expressed as H3cit-positive area per 100 neutrophils.

Quantification was performed blinded to condition using FIJI/ImageJ. Identical analysis parameters were applied to all images within each experiment. Data were pooled across fields for each biological replicate and presented as mean +/- SEM, with statistical tests specified in the figure legends.

### Kinetic analysis of SYTOX Green incorporation assay

Cells were plated at density of 5.10^4^ cells per well in Black/Clear 96-well Plates (Nunc) in Opti-MEM culture medium supplemented with SYTOX-Green dye (100ng/mL) (ThermoFisher, S7020) and infected/treated as mentioned in figure legends. Green fluorescence was measured in real-time using FLUOstar Omega plate reader (BMG Labtech) at 37°C with 5% CO_2_. Maximal cell death was determined with whole cell lysates from unstimulated cells incubated with 0.1% Triton X-100.

### Uronic acid measurement

Uronic acid (UA) was measured as described previously^12,56^. Kp strains were grown overnight in LB medium, at 37°C with constant agitation (200 rpm). The following day, cultures were diluted to an OD_600_ of 0.2 in M9 medium supplemented with 0.2% casamino-acids and incubated for 5 hours at 37°C, with constant agitation (200 rpm).

UA was extracted from 500µL culture with zwittergent, precipitated with ethanol and resuspended in tetraborate/sulfuric acid. Following addition of phenylphenol, UA was determined by absorbance at 520 nm. A standard curve was generated with glucuronic acid. Concentration was normalized to 1 OD_600_.

### Mucoviscosity assay

Mucoviscosity was quantified using a low-speed centrifugation assay, which assesses the ability of bacterial cultures to sediment under gentle centrifugal force, as described previously^12,56^. Kp strains were grown under the same conditions as described for the UA assay. After 5 hours of incubation, the OD_600_ of each culture was measured to determine total bacterial density. Cultures were then centrifuged at 1000xg for 5 min at RT. The OD_600_ of the supernatant were determined and plotted.

Strains exhibiting HMV resist sedimentation and therefore retain a higher proportion of bacteria suspended in the supernatant, resulting in higher post-centrifugation OD_600_ values. In contrast, non-HMV strains sediment efficiently, yielding lower supernatant OD_600_ readings.

### Transmission Electron Microscopy experiments

Samples were fixed using a final concentration of 2.5% glutaraldehyde and 2% paraformaldehyde in 0.1 M Sorensen buffer (pH 7.2; EMS, EMS11600-05), added to the culture medium in a volume-to-volume ratio for 15 minutes at room temperature. Following this primary fixation, the cells underwent secondary fixation with the same solution (2.5% glutaraldehyde, 2% paraformaldehyde in 0.1 M Sorensen buffer, pH 7.2) for 2 hours at room temperature and were then post-fixed at room temperature with 1% osmium tetroxide (OsO₄) and 1.5% potassium ferricyanide K3Fe(CN)6 in 0.1 M cacodylate buffer with 2 mM CaCl₂. Next, cells were treated overnight at 4°C with 2% aqueous uranyl acetate, dehydrated through a graded ethanol series, and embedded in Epon resin. After 48 hours of polymerization at 60°C, ultrathin sections (80 nm) were mounted onto 200-mesh Formvar-carbon-coated copper grids. Sections were then stained with Uranyless and lead citrate. Grids were examined using a transmission electron microscope (TEM; Jeol JEM-1400, JEOL Inc., Peabody, MA, USA) operated at 80 kV, and images were acquired with a digital camera (Gatan Orius, Gatan Inc., Pleasanton, CA, USA).

### Statistical analysis

All data are from at least three independent experiments. Statistical analyses were performed with Prism (GraphPad Software). All statistical tests and p-values are mentioned in the figure legends.

